# New gene evolution with subcellular expression patterns detected in PacBio-sequenced genomes of *Drosophila* genus

**DOI:** 10.1101/2022.11.30.518489

**Authors:** Chuan Dong, Li Zhang, Shengqian Xia, Dylan Sosa, Deanna Arsala, Manyuan Long

## Abstract

Previous studies described gene age distributions in the focal species of *Drosophila melanogaster*. Using third-generation PacBio technology to sequence *Drosophila* species we investigated gene age distribution in the two subgenera of *Drosophila*. Our work resulted in several discoveries. First, our data detected abundant new genes in entire *Drosophila* genus. Second, in analysis of subcellular expression, we found that new genes tend to secret into extracellular matrix and are involved in regulation, environmental adaption, and reproductive functions. We also found that extracellular localization for new genes provides a possible environment to promote their fast evolution. Third, old genes tend to be enriched in mitochondrion and the plasma membrane compared with young genes which may support the endosymbiotic theory that mitochondria originate from bacteria that once lived in primitive eukaryotic cells. Fourth, as gene age becomes older the subcellular compartments in which their products reside broadens suggesting that the evolution of new genes in subcellular location drives functional evolution and diversity in *Drosophila* species. Additionally, based on the analysis of RNA-Seq of two *D. melanogaster* populations, we determined a universal paradigm of “from specific to constitutive” expression pattern during the evolutionary process of new genes.

## Introduction

The understanding of gene origination and evolution over evolutionary time is central to understanding the genetic basis that shapes phenotypic and biological diversity. Novel genetic units arising recently at a specific genomic locus are regarded as new genes and are often taxonomically restricted to several species (Long et al. 2003; Ding et al. 2012; Chen et al. 2013). Such genes provide valuable materials for the study of the evolutionary questions and patterns because such genes maintain distinct characteristics compared with the old genes in terms of gene structure, epigenetic profiles, and transcriptional regulation (Werner et al. 2018). There are several known mechanisms that contribute to the creation of new genetic architecture such as: exon shuffling, gene duplication, retro-position, lateral gene transfer, and 7 other distinct molecular processes (Chen et al. 2013; Long et al. 2013). Among them, gene duplication is the most common molecular mechanism to generate new genes which is a widely observed phenomenon in all three domains of life (Zhang 2003). It is traditionally held that *de novo* origination of genes is not possible as every gene should emerge from pre-existing genetic materials (Jacob 1977; Ohno 2013), however our recent study showed that *de novo* gene origination is a common phenomenon in *Oryza spp.* (Zhang et al. 2019). Some researchers have even demonstrated that *de novo* genes emerging from proto-genes may be more prevalent than gene duplication in *Saccharomyces cerevisiae* species (Carvunis et al. 2012). These various mechanisms which lead to new gene origination perpetually occur throughout long periods of evolutionary time, thus the origination of new genes is important to explain biological diversity and genomic variation. New genes can rapidly become indispensable and acquire essential functions in diverse processes such as *de novo* genes related with human health (Toll-Riera et al. 2009; Van Oss and Carvunis 2019), new genes formed by duplication resolving intralocus sexual conflict in *D. melanogaster* (VanKuren and Long 2018; Schroeder et al. 2020), and new genes impacting reproductive behavior (Dai et al. 2008) and brain evolution (Chen et al. 2012). These phenomena are surprising as genes with critical biological functions are thought to be ancient and highly conserved. Later studies showed that new genes can reshape protein-protein and co-expression networks via gradually integrating into the ancestral interaction networks in genome-wide scale both for prokaryotes and eukaryotes (Zhang et al. 2015; Wei et al. 2016; Zu et al. 2019; Zhang et al. 2020). Such integration of new genes into pre-existing networks could promote the evolution of essential functions in new genes, causing them to become critical components in viability during development (Xia et al. 2016; Lee et al. 2019; Xia et al. 2020).

The *Drosophila* lineage has diverged into several species within the last 100 million years which provides excellent materials for studying new gene evolution and origination. Currently, three versions of gene age lists in *Drosophila* species have been published. Zhou *et*. *al*. published the initial gene age list for protein coding genes of *D. melanogaster* in 2008, which provided the first panoramic picture of the evolutionary origin of new genes in *D. melanogaster* (Zhou et al. 2008a), however they used only 5 outgroup species for age dating, greatly reducing the gene age resolution. In 2010 our group published the second analysis of gene ages for *D. melanogaster* in which we utilized 11 outgroup species to provide more detail about the origination of new genes (Zhang et al. 2010). Recently, Shao *et*. *al*. described an online knowledge of gene ages for several species, which is the most recent gene age inference for *D. melanogaster* (Shao et al. 2019). Our group re-sequenced several genomes of *Drosophila* species including *D*.*ananassae*, *D*.*erecta*, *D.hydei*, *D.novamexicana*, *D*.*persimilis,* and *D*.*virilis* with PacBio long-read sequencing technology. We also used PacBio to sequence *D.azteca* and *D.orena* for the first time as well as the most evolutionarily-related species to *Drosophila, Scaptodrosophila lebanonensis.* With the inclusion of *S*.*lebanonensis* we can investigate the process of gene birth at the most basal point of the *Drosophila* phylogeny which contains two subgenera, *Drosophila* and *Sophophora*, however previous studies on gene age dating in *Drosophila* species only focused on the *Sophophora* subgenus while the *Drosophila* subgenus has received little attention. In the present work we first used the new PacBio-based genome assemblies to conduct gene age dating in both for the *Drosophila* and *Sophophora* subgenera to produce a gene age list for *Drosophila* species. We used our new gene age list to study subcellular localization during gene evolution. Our results reveal several key insights. The large number of gene births identified may promote speciation and show new gene birth to be a common phenomenon both in *Drosophila* and *Sophophora* subgenera; Old genes tend to enrich in the mitochondrion, supporting the endosymbiotic theory while young genes tend to localize in extracellular space which may provide an environment for young genes that promotes their rapid evolution; As genes become older the number of subcellular compartments they localize to becomes broader, potentially driving functional and phenotypic evolution.

## Materials and Methods

### *Drosophila* PacBio sequencing

We utilized the PacBio long-read sequencing and assembly technologies to produce *de novo* or generate genome upgrade (GU) reference genomes for the set of phylogenetically informative species (Figure 1). This approach will create a genome-wide database that will make it possible to discover *Drosophila* genus-specific genes that will support the early divergence of the genus into two major subgenera, *Sophophora* and *Drosophila*.

**Figure 1.**
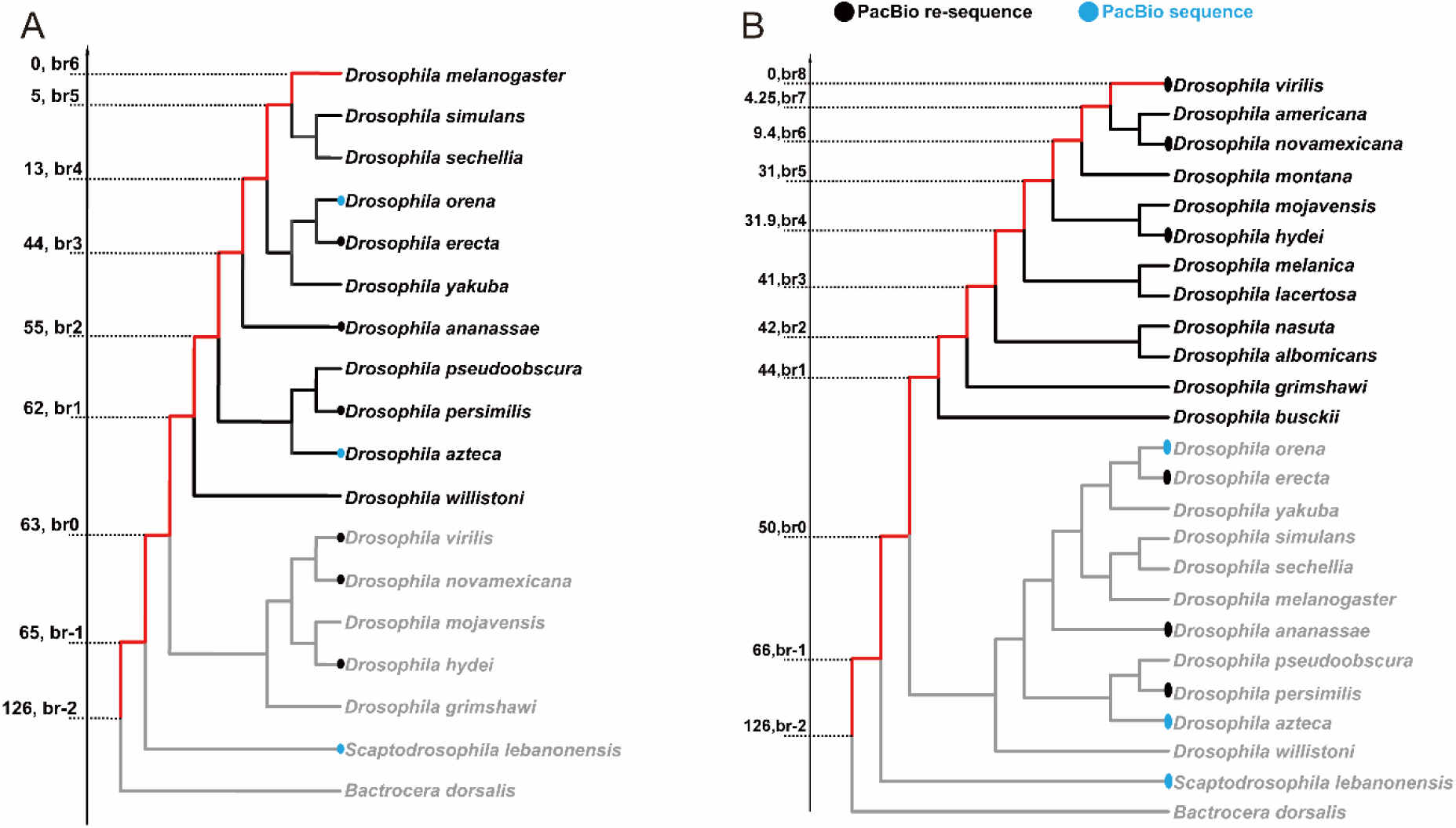
Evolutionary tree used for dating gene ages in *D. melanogaster* and *D. virilis*. A. Black circles indicate the re-sequenced species with PacBio long-read sequencing technology. Blue circles indicate species that were newly sequenced with PacBio. Species in black text are members of the *Sophophora* subgenus, the red line indicates the divergence direction of our focal species *D.melanogaster*, and those species in grey text are members of the *Drosophila* subgenus. B. Species in black text are members of the focal subgenus *Drosophila*, and the red line indicates the divergence direction of our focal species *D.virilis*, and those species in grey text are members of the *Sophophora* subgenus.

To determine the differences between the PacBio assemblies we created and previous assemblies used for detecting new genes (Zhang et al, 2010; Clark et al, 2007), we generated Mauve genome alignments of these assemblies relative to the same syntenic regions in *D. melanogaster* (Darling et al. 2004). Figure 2 provides clear evidence that the previous *D. simulans* assembly contains not only indel errors, but also sequence rearrangement errors, and draft assembly artifacts. In the case of unitig_14, the dark green, purple and pink blocks are significantly smaller, while the purple and pink blocks are placed in the incorrect positions (Figure 2A). In the case of unitig_9, the red block is significantly smaller in the available assembly, and yellow block is placed in the wrong position (Figure 2B), showing that long-read PacBio sequencing is capable of making significant improvements on existing *Drosophila* genome assemblies.

**Figure 2.**
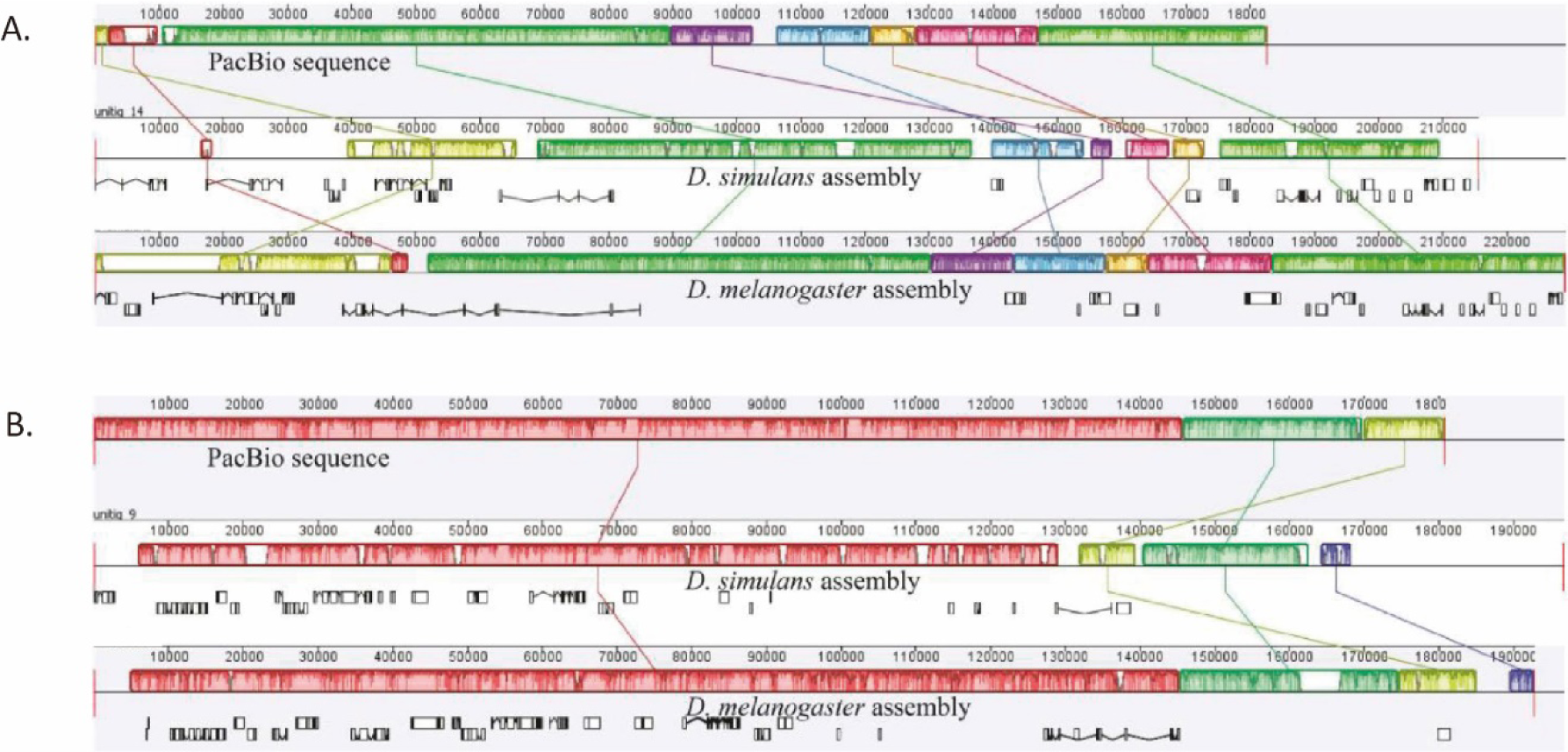
Alignment of two PacBio sequenced *D. melanogaster* BACs against their orthologous regions in the previous assemblies of *D. simulans* and *D. melanogaster*. Unitigs 14 and 9 are represented in panels A and B, respectively. Orthologous blocks are depicted with the same color. For each comparison, the PacBio sequence of *D. simulans* is on top, the *D. simulans* assembly is in the middle, and the *D. melanogaster* assembly is at the bottom. In both cases, PacBio sequences have exactly the same syntenic structure as the *D. melanogaster* assembly (BAC-based, high-quality), while the *D. simulans* assembly seems to be incorrectly assembled. Moreover, the *D. simulans* assembly has smaller ortholog blocks compared with the PacBio sequences and *D. melanogaster* assembly which further suggests that PacBio technology is also more accurate at the base pair level in reconstructing longer regions.

### Phylogenetic analysis and gene annotation of *Drosophila* species

The dating of gene ages depends on accurately inferred phylogenetic trees which should contain a focal species and several evolutionarily-related species. Focal species refers to the species which we want to conduct gene age dating of and the evolutionarily-related species are those species used as outgroups for whole genome alignment allow us to observe whether a gene is present or absent in outgroups. The TimeTree database (http://www.timetree.org/) provides species divergence times and accurate topological structures of species trees (Kumar et al. 2017a; Kumar et al. 2022). Therefore, to date the gene ages of *D. melanogaster* we obtained a phylogenetic tree containing our re-sequenced, newly sequenced, and previously sequenced species based on the TimeTree database (Kumar et al. 2017a). In total, we used 17 outgroup species for gene age dating of *D. melanogaster* resulting in 9 branches ranging from br-2 (branch -2) to br6 (Figure 1A and Supplementary File S1). All of the PacBio-based genome assemblies used in this work were created by us and are available for download from NCBI PRJNA475270 (https://www.ncbi.nlm.nih.gov/bioproject/PRJNA475270).

The *Drosophila* species contains two subgenera, *Drosophila* and *Sophophora*. Previous works on gene age dating for *Drosophila* spp. focused only on the *Sophophora* subgenus and paid very little attention to the *Drosophila* subgenus likely because the few species in the *Drosophila* subgenus in their reference tree. The evolution and origination of new genes in the *Drosophila* subgenus is an interesting topic which should be explored in greater details. We investigate gene birth using *Drosophila virilis*, a member of the *Drosophila* subgenus, as our focal species to outline the gene age distribution of this subgenus. There are merely 3 species in the *Drosophila* subgenus evolutionary tree that have been used to date gene ages of *D. melanogaster* (Zhang et al. 2010). Figure 1A is still insufficient for dating gene ages of *D.virilis* even though it contains our new PacBio-based assemblies and re-sequenced genomes (merely five species belong to *Drosophila* subgenus). To solve this problem, we collected genome assemblies for the 124 *Drosophila* species (Supplementary Table S1) available from NCBI at the time we performed this project.

We reconstructed a new phylogenetic tree focusing on the *Drosophila* subgenus using the following procedure. First, the 124 *Drosophila* species were compared with TimeTree. This comparison resulted in ruling out 18 *Drosophila* species from our initial species list because TimeTree did not have their records. Therefore, the initial tree reconstruction contains 106 leaf nodes (Supplementary File S2). Second, we uploaded the 106 *Drosophila* species to TimeTree so that we could reconstruct an evolutionary tree which we defined as the pre-tree. Depending on the pre-tree, we were able to judge which species belong to our focal subgenus. We selected species related to *D.virilis* in *Drosophila* subgenus from our initial tree to enrich the branches leading to *D.virilis*. Our selections were made based on the following two principles. First, we selected species belonging to the *Drosophila* subgenus because we needed to include more species to improve the gene age resolution in this subgenus. Second, the divergence time between our selected species and *D.virilis* needed to be within 100 million years. Species satisfying the above principles were selected. The final tree used to date gene ages in *D.virilis* ultimately contained 25 species and 11 branches (Figure 1B and Supplementary File S3). All of the genome assemblies used in dating the gene ages of *D. virilis* were downloaded from NCBI (Supplementary Table S2).

### Whole genome-based alignment

The traditional ways of identifying new genes is based on comparing percent identity of gene sequences and exon-intron structures. As gene duplication is a primary source of genetic innovation (Zhang et al. 2010), new genes derived from duplication would have similar sequences in reference species due to the sequence similarity between ancestral and duplicated genes. DNA duplication can lead to “many-to-one” or “many-to-many” orthologs in other species and would be treated as ancient genes when using traditional methods. Therefore, traditional gene dating methods have drawbacks and can underestimate the number of new genes. Zhang *et al* developed a synteny-based method to solve this problem (Zhang et al. 2010). Their method utilizes whole genome alignments to detect whether there is a syntenic ortholog in outgroup species for a specific gene. Most parts of our pipelines are following Zhang *et al*., 2010. Several whole genome alignments of the species considered here can be downloaded from the UCSC Genome Browser. Due to the lack of whole genome alignment data between *D.virilis* and its reference species as well as the newly the newly sequenced PacBio genomes (Figure 1B), we needed to perform the alignment by ourselves. We conducted whole genome alignment with LASTZ to obtain the alignments unavailable from the UCSC Genome Browser (Harris 2007). The parameters for LASTZ are following those used by UCSC Genome Browser: ‘O=400 E=30 K=2200 L=4000 H=2000 Y=3400’. MySQL and in-house Perl and Python scripts were used to store and analyze the data; and Linux shell scripts were further used to wrap the pipeline.

To obtain the present and absent information for a gene in *D. melanogaster* and *D. virilis* against their reference species showing in Figure 1A and Figure 1B, respectively, we mapped genes to the alignment between *D. melanogaster* or *D.virilis* and the outgroup species. Therefore, we downloaded annotations of *D. melanogaster* from Ensembl (version 97) and *D. virilis* from NCBI, respectively. We then compared gene annotations and genome alignments so that we could confirm the presence or absence of a gene in the reference species. If a gene had multiple transcripts due to alternative splicing, gene age was be determined by the oldest transcript.

### Subcellular compartment and the definition of enrichment index

COMPARTMENTS is an online resource which integrates subcellular localization information including data from manually curated literature, high-throughput screens, automatic text mining, and prediction from the “state-of-the-art” sequence-based prediction methods (Binder et al. 2014), therefore data from this resource is more reliable than these merely from bioinformatics prediction. In our work we downloaded subcellular location information from the COMPARTMENTS database. A confidence score ranging from 0 to 5 is conferred to quantify the degree of reliability for proteins located in a corresponding subcellular compartment. We performed kernel density estimation for the distribution of the confidence score obtained from the COMPARTMENTS database. Our result showed that most proteins have a confidence score approximately equal to 1, a relatively low confidence value (Supplementary Figure S1). To guarantee sufficient samples and the quality of subcellular compartments we only considered records with confidence score larger than 2, a relatively acceptable trade-off cutoff between confidence and the considered number of genes in our analysis. Additionally, we only considered 8 subcellular compartments including nucleus, cytosol, cytoskeleton, endoplasmic reticulum, golgi apparatus, plasma membrane, extracellular space, and mitochondrion. We used these subcellular annotation data to explore possible age-dependent subcellular localization relationships for proteins encoded by young and old genes in their evolutionary process. The gene IDs and their translated proteins were mapped with each other according to the gene annotation. For *D. virilis*, we employed DeepLoc 2.0 (Thumuluri et al. 2022) to predict the most reliable subcellular locations. There are ten possible cell compartments in its prediction result, in which plastid is a unique organelle in plant, we therefore remove this organelle in our analysis.

To quantify subcellular localization preference for a given protein and to remove the impact of background noise we proposed a parameter, EI (Enrichment Index), which measures the enrichment degree for proteins localizing in cellular compartments which can be calculated according to Equation 1.

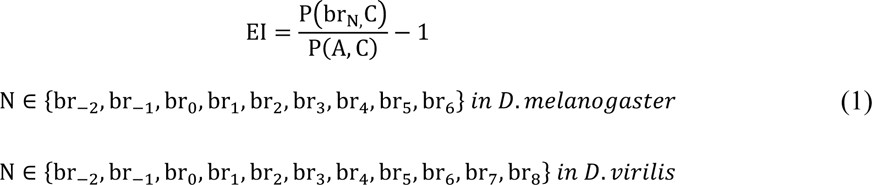

where P(br_N,_C) represents the frequency of genes in branch br_N_ whose translated proteins localize in a subcellular compartment C, and P(A, C) represents the frequency of all genes whose translated proteins localize in compartment C. There are 9 branches in our phylogenetic tree that used for dating *D*. *melanogaster* gene ages, and 11 branches for dating gene age in *D*. *virilis*. N ranges from the oldest branch br-2 to the youngest br6 in *D*.*melanogaster*, and for *D*.*virilis*, the N ranges from the oldest branch br-2 to the youngest br8. If the encoded proteins in a certain branch tend to enrich in a certain organelle, the frequency should be larger than the background frequency, i.e. P(br_N,_C) > P(A, C) and EI will larger than 0. If the encoded proteins are not enriched in a certain branch and a certain organelle, EI will be lower than 0. Finally, the subcellular location of proteins is mapped to their corresponding genes according to gene annotations of *D*. *melanogaster* and *D*. *virilis*, respectively.

## Results

### Large numbers of new genes identified from evolutionary lineages of two *Drosophila* subgenera

Genes ages ranging from br1 to br6 in *D. melanogaster* are defined as new genes (Zhang et al. 2010), and those genes from br1 to br8 in *D.virilis* are defined as new genes. According to these criteria we identified 846 young protein coding genes in *D. melanogaster,* which account for 6% (846/13890) out of all the dated protein-coding genes. Compared with previous work on gene age inference in *D. melanogaster*, the number of young gene estimation is more conservative (Zhang et al. 2010; Shao et al. 2019). According to these data, we estimate the birth rate of new protein-coding genes to be approximately 14 genes per million years (846/62) for the *Sophophora* subgenus, which presents an approximate equal gene birth rate (15 genes per million years) in contrast with our previous estimation (Zhang et al. 2010). Among the 846 young protein-coding genes there are 213 protein-coding genes distributing on br1, 218 on br2, 97 on br3, 236 on br4, 37 on br5, and 45 on br6 (Figure 3A and Supplementary Table S3_A), respectively.

**Figure 3.**
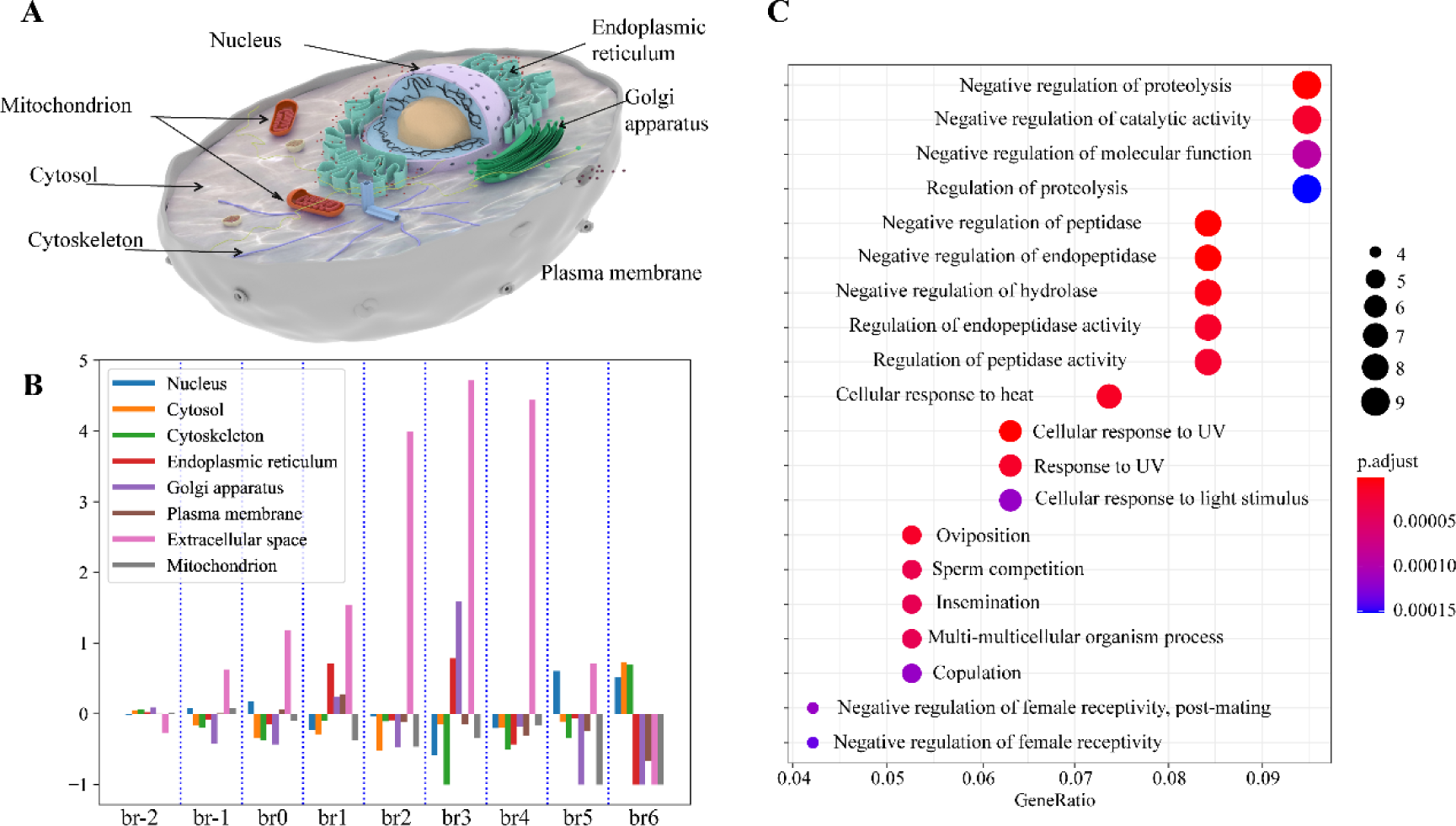
Gene age distribution and age comparison with GenTree. A. age distribution of protein-coding genes for *D. melanogaster*; B. age distribution of ncRNA genes for *D. melanogaster*; C. age distribution of protein-coding genes for *D. virilis*; D. age distribution of lncRNA genes for *D. virilis*; E. gene age comparison between our results and GenTree, where red values on the left indicate branches in Shao *et al*’s work. “→” means the gene age was changed in our work compared with Shao *et al*’s work. The number in parenthesis indicates how many genes were moved from Shao *et al*’s branch to our newly dated gene age list. For example, “br0→br-2, br-1 (11399)” means there are 11399 genes in branch br0 in Shao *et al*, however, those genes were changed to br-2 or br-1 in our work. “✓” means that if we removed the newly added PacBio species, the gene age will be same in the two gene age version.

Remarkably, the introduction of *S. lebonenisis* and *B. dorsalis* led us to identify 943 protein-coding genes that predated the divergence of the two subgenera (*Drosophila* subgenus and *Sophophora* subgenus) in the focal species *D. melanogaster*. Similarly, in the focal species *D. virilis*, we also identified 778 protein-coding genes that appeared before the divergence of the two subgenera, which contained 333 gene pairs sharing sequence similarity (detected by a PSIBLAST search at an e-value of 10e-3). These high numbers of new genes were created in only ∼1-6 million years of the common ancestral species (CAS) of *Drosophila* genus, revealing an unusually high origination rate of new genes, which can be as high as 943 per million years as reflected as the *D. melanogaster* focal species, associated with the speciation and short evolutionary period of CAS before its divergence to two major subgenera.

In the focal species of *D. melanogaster*, we also obtained 322 young ncRNA genes (only refer to the ncRNA annotated in Ensembl database) representing 13.03% (322/2470) of the all annotated ncRNA genes (Supplementary Table S3_B). We estimated the young ncRNA gene birth rate to be 5 genes per million years after the divergence from the *Drosophila* subgenus. Among those young ncRNA gene there are 90, 107, 20, 86, 13, and 6 ncRNA genes distributed on br1 through br6, respectively (Figure 3B and Supplementary Table S3_B). The introduction of *S. lebonenisis* and *B. dorsalis* also led to the identification of 702 *Drosophila* subgenus specific non-coding genes that predated the divergence of the two subgenera (*Drosophila* and *Sophophora*). Based on our observations, there is also a large number of protein-coding gene and ncRNA gene births when forming *Drosophila* species (br-1), indicating that gene birth might be an important driver of speciation and phenotypic evolution.

To test this hypothesis, we enriched our *Drosophila* subgenus phylogeny by adding more species (Supplementary Table S2), resulting in a phylogeny with 25 outgroup species and 11 branches (from br-2 to br8). With this more refined evolutionary tree we annotated the gene age of *D. virilis*, and the same phenomenon was found, that is a large number of gene births when forming *Drosophila* species (Figure 3C, Figure 3D and Supplementary Table S4) for both protein-coding and long non-coding RNA (lncRNA) genes which might further indicate that gene birth drives speciation. Among the dated genes in *D. virilis* 671 protein-coding genes were inferred to be young genes which account for 5% (671/13648) among all dated protein-coding genes. Therefore, the rate of gene birth for protein-coding genes in the *Drosophila* subgenus is estimated to be 17 genes per million years (671/38) which is nearly the same rate of protein-coding gene birth in the *Sophophora* subgenus. There are 642 lncRNA genes annotated as young genes (38% among the dated lncRNAs of *D.virilis*) resulting in the lncRNA birth rate in the *Sophophora* subgenus to be 16 (642/38) genes per million years. Although the birth rate of protein-coding gene is nearly same among the two subgenera, the birth rate for the lncRNA genes in the two subgenera is distinct, which imply that the formation of new regulatory network might be more important than the birth of protein-coding gene itself during species formation.

A comparison with previous databases in the *Sophophora* subgenus (Zhang et al, 2010; Shao et al, 2019) that we created previously with different sequencing and annotations reveals improvement by the third-generation sequencing and annotation. In the earlier database, we found that except br2 all other branches are identified with fewer new genes in this study. The higher quality genomes created by the long-read PacBio detected more homologous genes in related species (Materials and Method), leading to changes in dating a small portion of genes.

We also compared the dating results in this study with those reported in GenTree (Shao et al, 2019). We found that 7.9% of genes are not annotated either in our databases or the GenTree database: 148 protein-coding genes not in our annotation while 955 protein coding genes we identified are have no annotations in GenTree. (Figure 3E). Further age comparison of the overlapping genes (12935) revealed 1047 genes annotated with the same age whereas 11588 genes present the older gene age compared with GenTree, which is due to our inclusion of two reference species outside *Drosophila*, especially increasing branches br-2 and br-1. 11399 out of the 11588 genes on br0 in GenTree were moved to br-2 and br-1 in our database. If we removed br-2 and br-1 in our phylogenetic reconstruction those genes would indeed correspond to br0 (Figure 3E). Similarly, after removing all the newly added species, the gene age distribution of 11588 genes (11366+82+34+38+51+8+9) closely resembled the branch placements in GenTree, with approximately 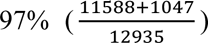 having the same branches placement, illustrating the high consistency of the two gene age versions with approximately 3% genes having different ages between the two databases. This emphasizes the importance adding high quality PacBio-based assemblies and more outgroup species to improve the resolution and accuracy of gene age inference.

Overall, the distributions of the new genes in the two backbone subgenera, *Drosophila* and *Sophophora,* present high numbers of evolutionary new genes with similarly rapid origination patterns across phylogenetic branches in the both lineages. Surprisingly, the short period of speciation and evolution of the common ancestral species before the genus diverged was associated with a high rate of new gene evolution, suggesting a common and extensive requirement of newly evolved *Drosophila* genus for new gene functions at protein and RNA levels in the newly occupied ecological niches of the genus.

### Subcellular location analyses of *D. melanogaster* revealed enrichment of young genes in extracellular expression

In the analysis of *D. melanogaster*, we considered 8 subcellular compartments, including nucleus, cytosol, cytoskeleton, endoplasmic reticulum, golgi apparatus, plasma membrane, extracellular space, and mitochondrion to explore age-dependent subcellular localization for proteins encoded by young and old genes (Figure 4A).

**Figure 4.**
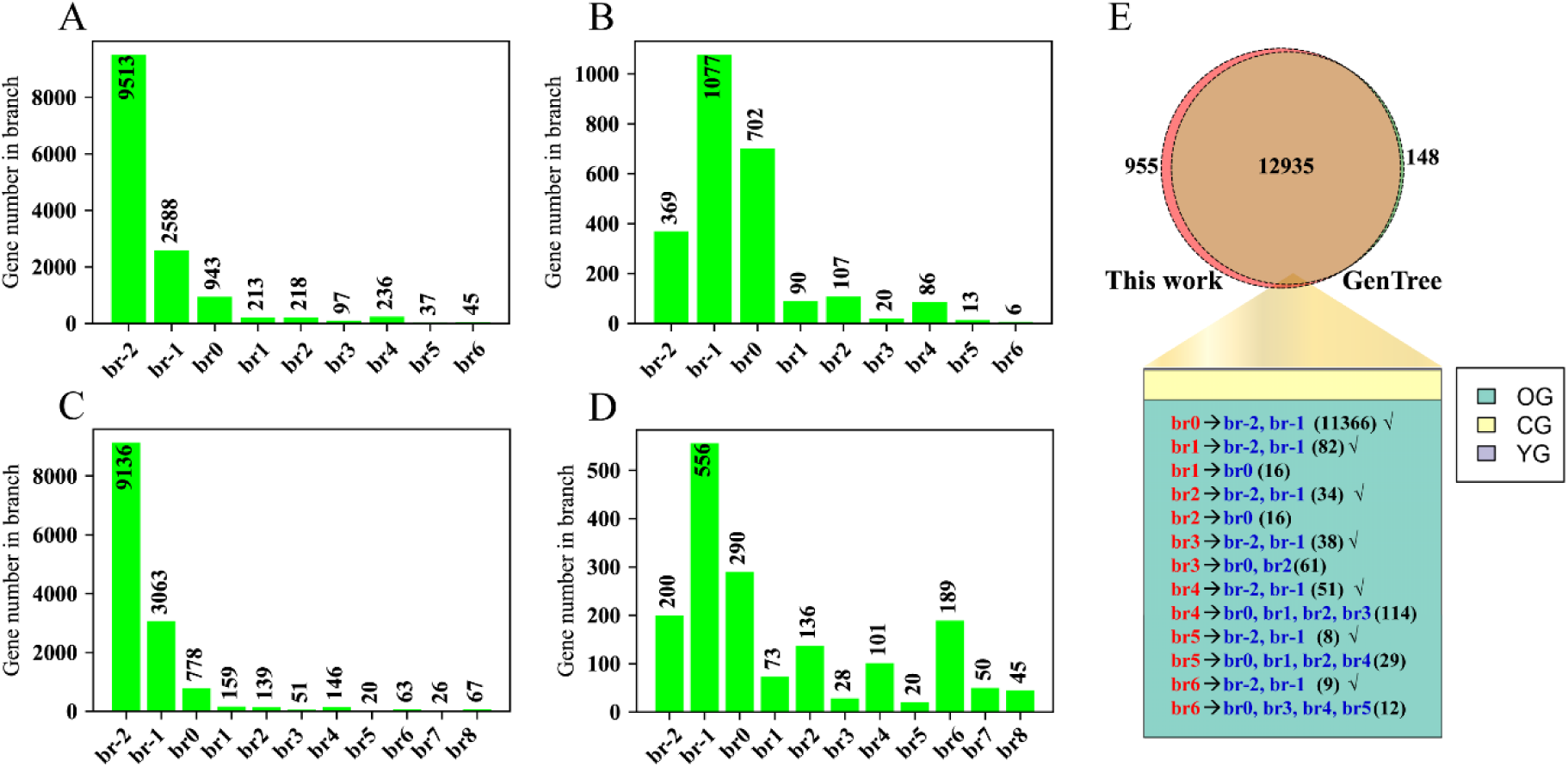
Enrichment analysis in subcellular compartments for proteins encoded by different gene ages. A. An illustration of subcellular compartments with those used in this study highlighted; B. EI index v.s. gene age analysis; C. gene ontology analysis for young genes whose proteins can export out extracellular space.

We calculated EI for all the genes in each branch and each of the 8 compartments that we considered in this work relying on equation 1 (method part). According to our EI, we found that young protein-coding genes tend to secrete their products outside of the cell and become components of extracellular space, especially for the genes distributed among branches br2, br3 and br4 (Figure 4B). Why do those young gene encoded proteins secrete outside of the cell, and what are the functions of their proteins located in extracellular space? To investigate this we performed gene ontology analysis using clusterProfiler (Yu et al. 2012; Wu et al. 2021) and org.Dm.eg.db annotation packages (Carlson 2013). Our results indicated that genes secreting into extracellular space can be divided into three functional categories: negative regulation of enzyme activity for peptidases and endopeptidases, environmental adaption (especially for the function of heat and light reaction), and functions related to reproduction (Figure 4C). The phenomenon of proteins encoded by young genes being involved in environmental adaption supports the hypothesis that new genes promote species-specific genetic novelties which could impact their fitness and contribute to biological diversity. For example, a recent study found that a novel chimeric gene family originating 48–54 million years ago promotes adaptive evolution of *Danioninae* fishes (Fang et al. 2021; Long 2021). Other studies have also reported that young genes often have relatively high and specific expression patterns in testis (Carelli et al. 2016; Guschanski et al. 2017), highlighting the functional relationship between young genes and species-specific reproduction. The analyses of gene expression from previous works and the ontology analysis in our study both support our hypothesis that young genes play critical roles in species-specific reproduction.

A previous study showed that the subcellular location of proteins in mammalian species can affect their evolutionary rate, and that genes whose proteins are exported out of the cell often evolve faster than those whose proteins localized within cells (Julenius and Pedersen 2006). The proteins encoded by young genes show the hallmark of tending to secrete extracellular space, promoting these proteins can interact with the extracellular environment directly, meanwhile these proteins can be easily impacted by environmental fluctuations. We, therefore, infer that extracellular space can provide an amenable environment for the fast evolution of proteins encoded by young genes, thus promoting functional and phenotype novelty.

### Enrichment of genes in mitochondrion localization with evolutionary ages in *D. melanogaster*

The above analysis raised very intriguing questions regarding how EI (enrichment index) change in the subcellular localization between young and old genes during their evolution. The investigation of this question will be valuable in understanding the relationship between functional novelties, phenotype diversities, and protein subcellular localization. We divided genes from *D. melanogaster* into 9 levels (br-2 to br6) according to their age and observe how the EI index changed over evolutionary time. We calculated the Pearson correlation coefficient between the 9 different age levels and EIs among all of the 8 subcellular compartments. To calculate this a series of values ranging from -2 to 6 were conferred to protein-coding genes to represent their age within the *Drosophila* phylogeny. We observed that proteins encoded by old genes located in the plasma membrane and mitochondrion have a relatively higher EI in contrast with proteins encoded by young genes, and that EI increases as gene age becomes older, i.e. *r*=-0.783, *p-value*=0.0126 for plasma membrane and *r*=-0.848, *p-value*=0.0039 for mitochondrion (Figure 5). Labelling the gene branch levels by numbers cannot reflect real divergence times, thus we utilized the divergence time obtained from TimeTree (Kumar et al. 2017b) to repeat the above analysis. We obtained the same strong and significant correlation between the EI and the real divergence time with respect to plasma membrane (r=-0.696, p-value=0.0375, Supplementary Figure S2) and mitochondrion (r=-0.729, p-value=0.0258, Supplementary Figure S2). The Spearman’s rank correlation coefficient also represents the same trend for proteins localized in mitochondrion and plasma membrane. The mitochondrial endosymbiotic theory asserts that mitochondria originated from a bacteria that once lived within primitive eukaryotic cells (Sagan 1967). According to this theory ancient genes should be observed in mitochondria. As expected, we found that old genes to have relatively larger EI for mitochondrion compared with young genes which supports the mitochondrial endosymbiosis theory. Therefore, our conclusion is consistent with the inference of the endosymbiotic theory and provides insight into endosymbiotic theory from another aspect.

**Figure 5.**
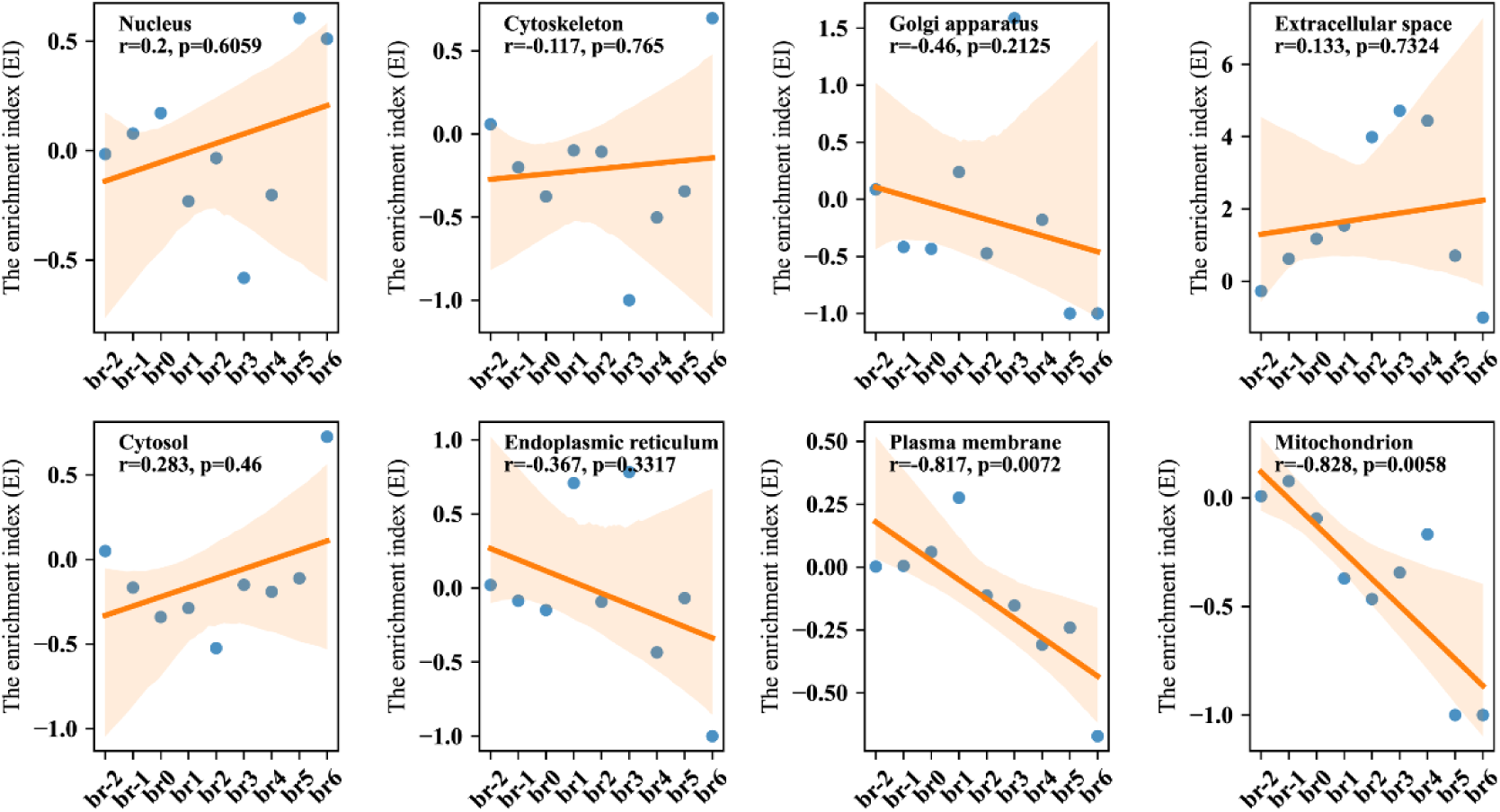
Correlation analysis between gene age and EI among the eight subcellular compartments based on *D. melanogaster*. The orange lines are the fitted line between gene age and EI index. The light orange background represents the 95% confidence interval.

### New genes of *D. melanogaster* created an increasing diversity of protein subcellular localizations in evolution

Does the number of subcellular locations for proteins encoded by genes change over evolutionary process? To explore this, we divided our gene list in *D. melanogaster* into four catalogs according to age which include genes in br-2 (the oldest group), genes in br-1 (the older group), genes in br0 (old group), and genes distributed in br1 through br6 (young group). One gene may encode several proteins due to alternative splicing, therefore multiple subcellular compartments from multiple proteins can be simultaneously mapped to one gene. We observed that 15.1031% of genes in br-2 localized in more than three subcellular compartments and that the proportion of genes located in more than three subcellular compartments decreased as gene age decreased. For example, we observed that only 6.1091%, 5.3691% and 3.8997% of genes on br-1, br0, and the young gene catalog that can be localized in more than three subcellular compartments (Figure 6A).

**Figure 6.**
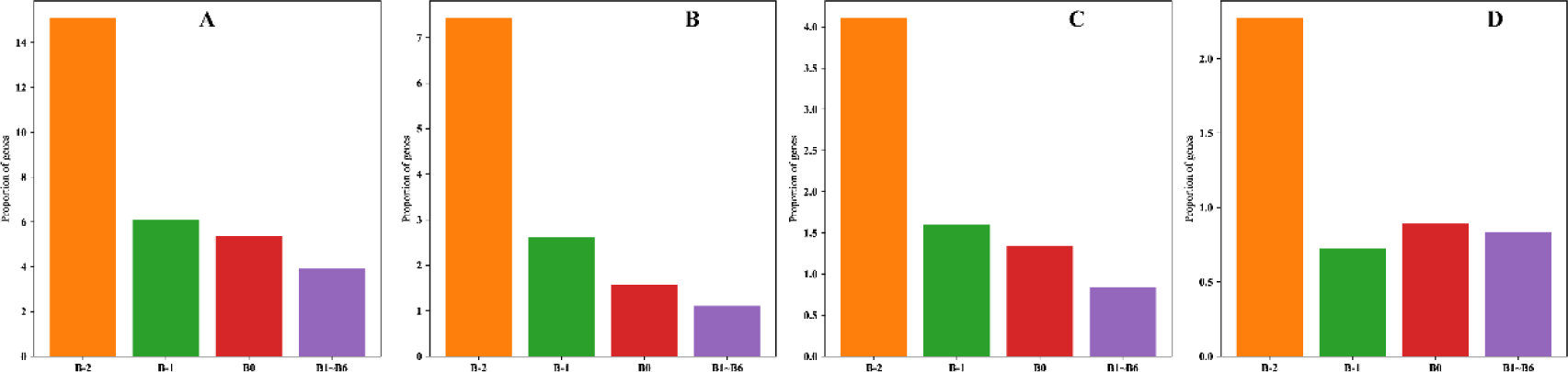
The proportion of subcellular location among different gene age catalogs based on *D. melanogaster*. X-axis represents gene categories divided by gene age. Y-axis represents gene proporation at a certain threshold, which means the proportion of gene having the number of subcellular compartments larger than this threshold.

Based on the same statistical pipeline we calculated the proportion of genes localizing in more than 4, 5, or 6 compartments. Our results illustrate that proteins encoded by old genes tend to localize in a wider range of compartments than proteins encoded by young genes according to the percentage of genes located in the number of more than 4 (Figure 6B), 5 (Figure 6C) and 6 (Figure 6D) subcellular compartments. Together, these results at different cutoffs provide solid support for our conclusion that the range of localized organelles becomes broader as genes become older.

Previous evidence demonstrates that the change of subcellular location of a protein or lncRNA can strongly influence and alter protein functions (Julenius and Pedersen 2006; Simpson and Pepperkok 2006; Byun-McKay and Geeta 2007). Guo et al recently report a pair of lncRNA orthologs in human and mouse which show different subcellular localization which contribute to their functional evolution (Guo et al. 2020).

Ashley et al proposed that subcellular re-localization can promote new gene birth because the change of one amino acid in signal peptides can influence subcellular location (Byun-McKay and Geeta 2007). Based on the analysis, we propose a hypothesis to explain how the subcellular locations of proteins encoded by genes affect functional diversity. First, over time, genes acquire more variable splicing forms, producing more encoded proteins, each carrying different subcellular localization signals to guide their localization with different subcellular compartments; Second, the highly refined subcellular compartments of higher eukaryotic species divide different proteins encoded by the same gene, creating different subcellular environments; Third, different subcellular compartments exert different selection pressures on their respective proteins and thus promote functional diversity. From the perspective of protein localization, the function diversity during gene evolution can be can be description by: (1) from the diversity of alternative splicing to diversity of subcellular location, and diversity of cellular compartments create different environmental selection pressures, which in turn act on gene products to promote functional and morphological diversity.

### The consistent subcellular localization pattern is also present in *D. virilis*

We employed DeepLoc2.0 (the accuracy prediction model was activated by -m option) to predict subcellular locations based on the protein list in *D.virilis* followed by mapping the proteins to their corresponding protein coding genes according to the gene annotation. DeepLoc2.0 can classify proteins to 10 subcellular compartments (cytoplasm, nucleus, extracellular, cell membrane, mitochondrion, plastid, endoplasmic reticulum, lysosome/vacuole, golgi apparatus, peroxisome) (Thumuluri et al. 2022), among which plastid is a unique organelle in plant. therefore we removed this organelle in our analysis. We also found that old genes tend to localize in the mitochondrion compared with young genes (r=-0.788, p=0.004), which demonstrate the similar pattern for new gene subcellular location in mitochondrion both for *Drosophila* and *Sophophora* (Figure 7A).

**Figure 7.**
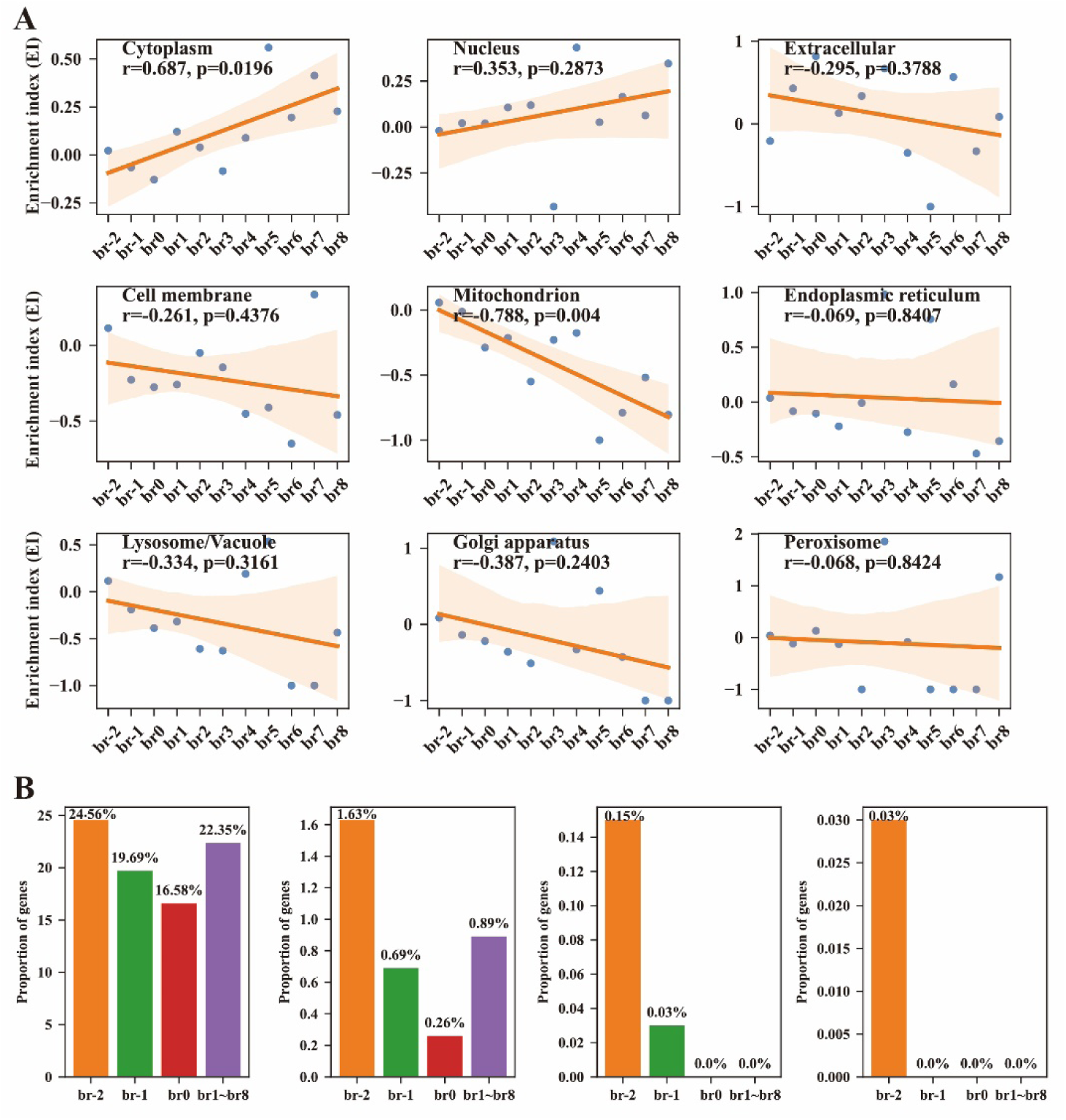
The subcellular location pattern based on *D.virilis*. A. EI index v.s. gene age analysis based on *D.virilis*; B. The proportion of subcellular location among different gene age clusters based on *D.virilis*.

We also explored the number of protein coding genes of different subcellular localizations during their evolutionary process to answer that if this pattern is still maintained in *D. virilis* that we ever found *in D. melanogaster*. Four gene age catalogs were created, in which br-2 is the oldest branch following by br-1, br0 and the young group from br1 to br8. The gene percentages localized in more than four different cutoffs among each catalog suggests that the gene proportion in the oldest branch (br-2) always larger than these in young group (Figure 7B). When we set the cutoffs as 4 and 5 (gene percentage above 4 and 5 cellular compartments), the gene percentage in the young group was reduced to 0. Additionally, as gene becomes older, the percentage of gene number above a certain cutoff gradually increase from br1, br0, br-1 to br-2 except for genes in the young group (Figure 7B).

The young protein-coding genes usually have a weak subcellular localization signal, thus can introduce noises for this analysis in *D. virilis* analysis, which might be the reason for the unexpected gene percentage in the young group (br1∼br8). However, for such group at the cutoffs above 4 and 5 compartments, we did not detect any genes which can be localized in more than such number of subcellular compartments. Conclusively, the subcellular location in mitochondrion and the number of localized subcellular compartments during the evolution of young genes show the same pattern both for *D. melanogaster* and *D.virilis*.

### The age patterns of gene expression reveal a dynamic process of directional gene traffics between the X chromosomes and autosomes

We defined these genes as sex-biased gene if these genes exhibit differential and significant expression patterns between the two sexes of *D. melanogaster*. We considered a gene to be a male-biased gene if the gene’s expression pattern can be characterized by equation (2), where male means the gene expression in male and the counterpart female means the gene expression in female. Similarly, a gene is considered as female-biased genes if this gene’s expression pattern can be characterized by equation (3).

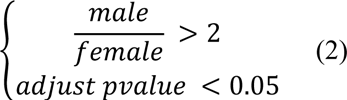

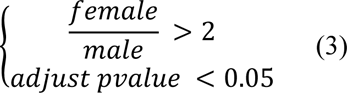

To calculate the sex-biased expression genes, we downloaded the raw RNA-Seq data (GSE99574) associated with *D. melanogaster* from NCBI https://www.ncbi.nlm.nih.gov/geo/query/acc.cgi?acc=GSE99574, which was provided by Yang et al and Mahadevaraju et al (Yang et al. 2018; Mahadevaraju et al. 2021). Several programs were employed to finish this task, in which the fastp (version 0.23.2) was utilized to quality control the raw RNA-seq data (Chen et al. 2018), then hisat2 (version 2.2.1) was employed to map sequencing reads to *D. melanogaster* genomes (Kim et al. 2019) with utilizing samtools (version 1.14) to transform the mapping file format (Danecek et al. 2021), followed by featureCounts (version 2.0) and DESeq2 (version 1.30.1) to count read number for each gene and to compute differential expression genes (Liao et al. 2014; Love et al. 2014).

GSE99574 includes two *D. melanogaster* populations, w1118 and orgR, therefore we have three sex-biased gene sets calculated by the two populations: sexed-biased genes detected according to w1118 (Supplementary Table S5), sexed-biased genes detected according orgR (Supplementary Table S6), and the intersected sexed-biased genes between w1118 and orgR. According to gonad tissue of *D. melanogaster*, we observed that:1) Young male-biased genes are enriched in the X chromosome; 2) Old male-biased genes are scarce (ranging from 29% to 31%) in X chromosome compared with young male-biased genes (ranging from 77% to 86%) in X chromosome; 3) There is no difference for old sex-biased genes on X chromosome; 4). Young female-biased genes are scarce both in the X chromosome and autosomes, however old female-biased genes are enriched in the X chromosomes and autosomes compared with young female-biased genes; 5) Old female-biased genes are slightly enriched in the X chromosome compared with old male-biased genes. Therefore, we concluded that X chromosome shows a paucity of old male-biased genes (Figure 8A, Figure 8C and Figure 8E), however young-male biased genes are enriched in X chromosome (Figure 8B, Figure 8D and Figure 8F). These observations are holded among our three sex-biased gene sets, which is consistent with our previous work(Zhang et al. 2010). We conclude that such a distribution is not a random phenomen for male-biased genes, but is a real pattern. We also utilized the whole body expression data from the two *D. melanogaster* populations to repeat the above analysis and again found that the X chromosome shows a paucity of old male-biased genes and an excess of young male-biased genes (Supplementary Figure S3, Supplementary Table S7 and Supplementary Table S8). To verify the reliability of the sex-biased genes, we performed correlation analysis of the intersected sex-biased genes from the two populations according to their expression fold change. The results showed that the correlation approaches more than 0.9 both for gonad (Supplementary Figure S4) and whole body (Supplementary Figure S5). That is to say the log2(fold change) of our sex-biased genes from two independent populations (w1118 and orgR) of the same species *D. melanogaster* have strong consistency. Therefore, we can conclude that the sex-biased genes are reliable and our conclusions based on gonad and whole-body data from the two population is solid and reliable.

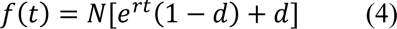

**Figure 8.**
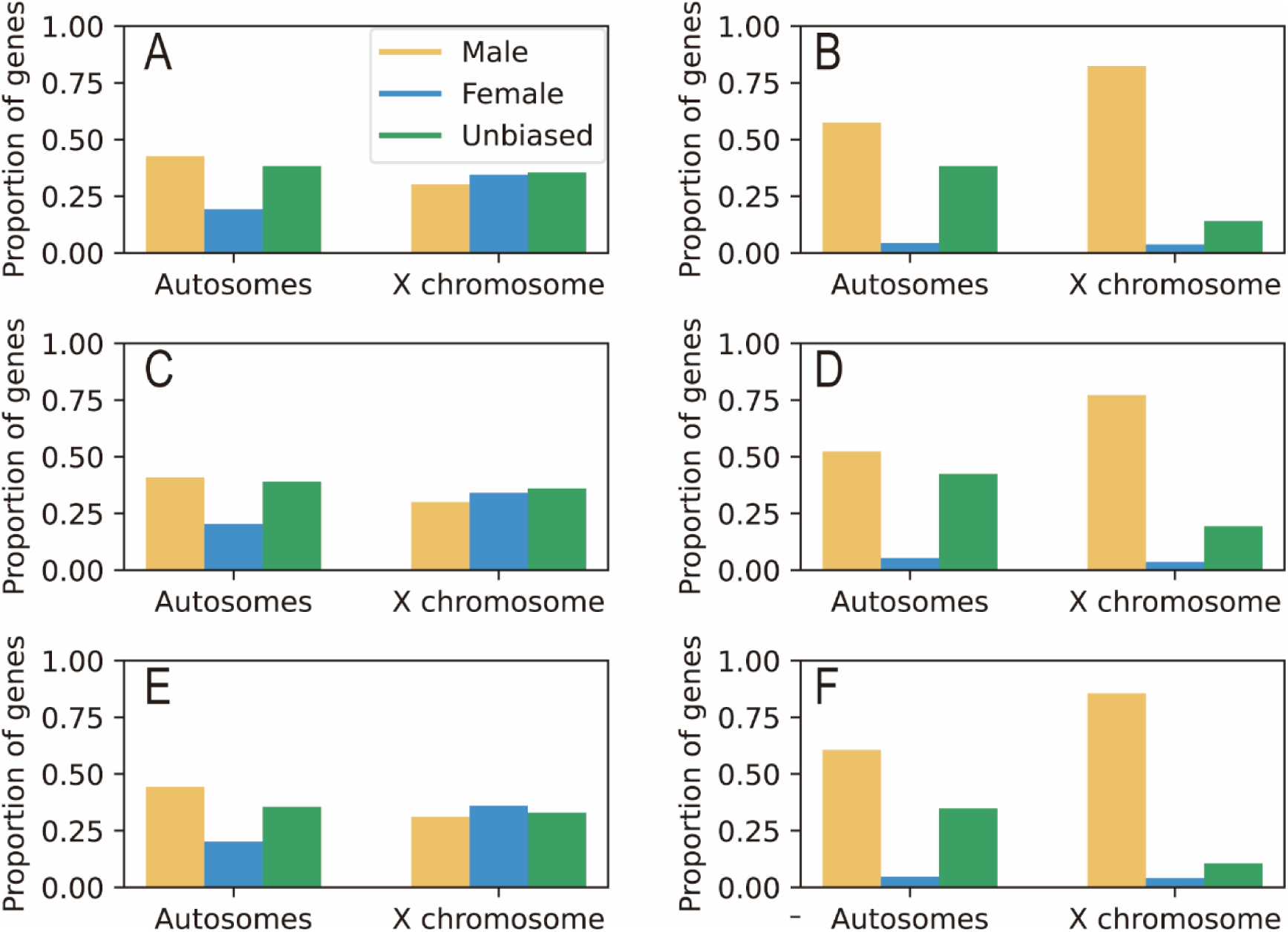
The distribution of male-biased genes among young and old gens. A. the distribution of sex-biased and unbiased old genes (based on w1118) and; B. the distribution of sex-biased and unbiased young genes (based on w1118); C. the distribution of sex-biased and unbiased old genes (based on orgR) and D. the distribution of sex-biased and unbiased young genes (based on orgR); E. the distribution of sex-biased and unbiased old genes (based on intersected parts of w1118 and orgR) and F. the distribution of sex-biased and unbiased young genes (based on intersected parts of w1118 and orgR).

Based on our new gene age data and expression in gonad from W1118 and orgR populations, we modeled the genomic distribution of male-biased genes between autosomes and the X chromosome over evolutionary time according to our previous used exponential decay formula (equation 4) (Zhang et al. 2010). A larger proportion of recently evolved male-biased genes, such as these young genes created in br5 (within in 13 Myr) and br6 (within 5 Myr), were enriched in the X chromosome (For example, 86.7% young genes created in br5 and br6 are male-biased genes based on our analysis of gonad tissue from W1118, and 85. 7% young genes created in br5 and br6 are male-biased genes based on the analysis of gonad tissue from orgR). The combined data from gonad tissue of W1118 and orgR also presented the same pattern, which illustrates a distinctive pattern for those genes originating on older branches, especially for those genes in br-2 (10% in w1118 gonad tissue, 11.1% in orgR gonad tissue, 10.8% in the combined gonad of w1118 and orgR populations) and br-1 (12.8% in w1118 gonad tissue, 12.7% in orgR gonad tissue, 13.2% in the combined gonad of w1118 and orgR populations). These data clearly show a decreased ratio compared with the genomic background of X-linked genes (∼17%).

To avoid observation bias caused by gene expression fluctuation in gonad tissue of w1118 and orgR populations, we used the gene expression data from whole body to re-model this process. We also observed that the recently created male-biased young genes, especially for genes in br5 and br6, are enriched in the X chromosome, but old male-biased genes are scarce in the X chromosome (Supplementary Figure S6). Therefore, our analysis combining two tissues (gonad and whole body) and two populations (w1118 and orgR) strongly support our observed expression pattern for young male-biased genes during their evolution.

These observations immediately suggested a directional enrichment in evolution of sex-specific expression between the X chromosomes and autosomes. Taking advantage of the more outgroup species genomes than previous database (Zhang et al, 2010. Genome Res; Clark et al, 2007. Nature), we analyzed the sex-biased expression and chromosomal locations of genes in different evolutionary branches. The proportions of male- and female-biased genes reveal a gradual shifting pattern between the two types of chromosomes with the ages of genes (Figure 9). The three sets of data all showed that the most recent young genes (branch 6) contributed 85% of sex-biased genes for male-biased expression and 15% for female-biased expression, then dropped to 23% for male-biased expression and 77% for female expression when gene ages reached 28 MYA. When gene ages approached 50-63 MYA, the proportion further decreased with fluctuation to 15-32% (average 20%) for male-biased expression and correspondingly 85-68% for female expression. The three old or ancient branches (br0, br-1 and br-2) reached to a stable frequency around 13% for male biased expression whereas 87% for female expression on X chromosome.

**Figure 9.**
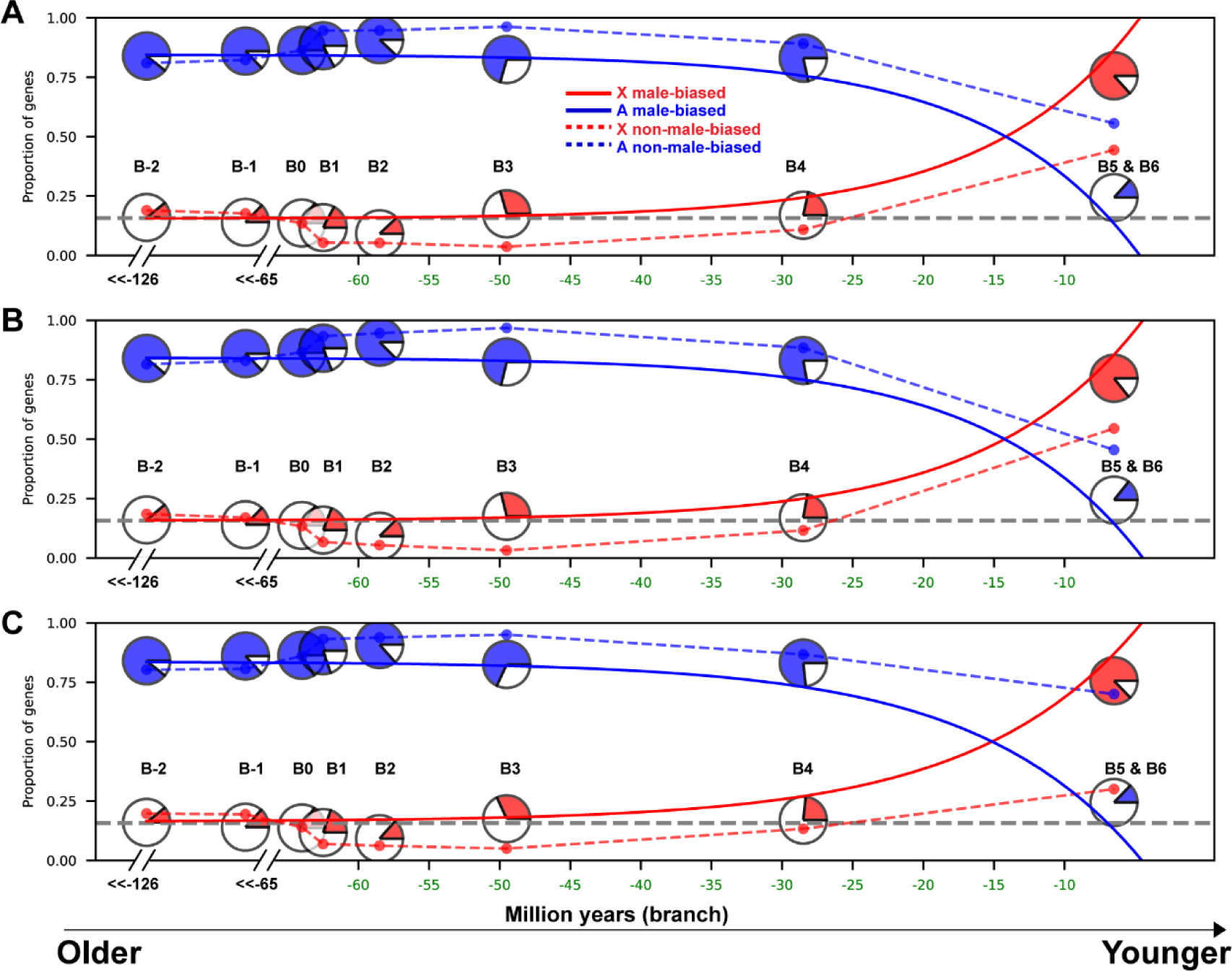
The modelling of male-biased gens in evolutionary course. A. Proportions of male-biased and non-male-biased (female-biased and unbiased combined) genes originating in different evolutionary periods based on w1118. To model the origination process of male-biased genes, which converged as: N = 1.466, r = 0.0943, and d = 0.105. B. Proportions of male-biased and non-male-biased (female-biased and unbiased combined) genes originating in different evolutionary periods based on orgR. To model the origination process of male-biased genes, which converged as: N = 1.425, r = 0.0917, and d = 0.11. C. Proportions of male-biased and non-male-biased (female-biased and unbiased combined) genes originating in different evolutionary periods based on the intersected parts of w1118 and orgR. To model the origination process of male-biased genes, which converged as: N = 1.396, r = 0.0858, and d = 0.116.

### The age-dependent pattern of gene expression reveals a “from specific to constitutive” paradigm

We downloaded the raw RNA-Seq data from NCBI (GSE99574), which includes 7 tissues of *D. melanogaster* followed by calculating the gene expression by our self-written pipeline (https://github.com/RiversDong/DEGs) by integrating hisat2 (Kim et al. 2019), StringTie (Pertea et al. 2015; Shumate et al. 2022) and fastp (Chen et al. 2018). Each tissue in GSE99574 contains 4 biological replicates, and we performed correlation analysis for each pair of biological replicates of a certain tissue in male and female from the two populations (w1118 and orgR). A high correlation coefficient indicates better repeatability, while the opposite indicates that there may be a problem with a particular biological replicate. Our results showed that the gene expression among each pair of biological replications shows high correlation except for a biological replication of genitalia in male from w1118 population (from Figure 10A to Figure 10G). For this replicate, the correlation of sample 1 with other replicates presents relative low relevance, however the other pairs show high correlation (more than 0.95) indicating that gene expression calculated by sample 1 is not correct (Figure 10E). We therefore excluded this sample from our analysis in this part.

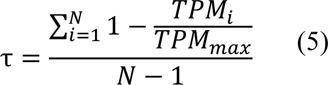

**Figure 10.**
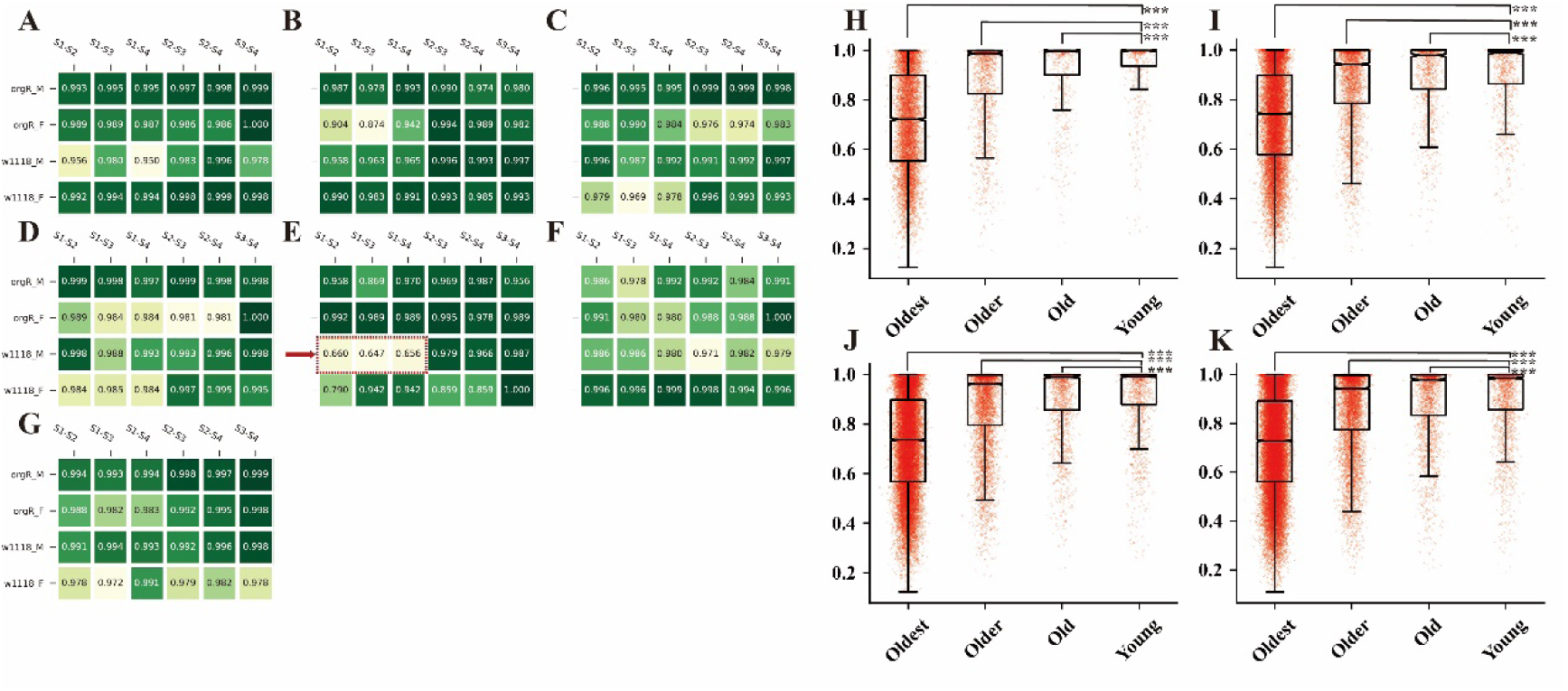
The expression pattern of new genes in multi-tissues. A to G illustrate the coefficients of Pearson correlation analysis between each pair of biological replicates. A. reproductive system without gonad and genitalia; B. thorax without digestive system; C. head; D. Gonad; E. genitalia; F. digestive plus excretory system; G. abdomen without digestive or reproductive system. H. gene expression pattern cross tissues in male of orgR population; I. gene expression pattern cross tissues in female of orgR; J. gene expression pattern cross tissues in male of w1118; K. gene expression pattern cross tissues in female of w1118; *** denotes significant defiance detected by Wilcoxon rank-sum statistic test.

Yanai et al (Yanai et al. 2005) proposed τ index to evaluate the tissue specificity expression for a gene. The τ index is range from 0 to 1 reflecting how specific tissue expression for a gene. More specifically, a high τ value illustrates a narrow expression across tissues, however a low τ value illustrates a border expression across tissues. We calculated the tissue specificity index according to equation 5 and the expression data calculated from 7 tissues of two *D. melanogaster* populations after excluding the first sequencing genitalia sample from w1118 male. Accordingly, we found that young genes show a narrow and significant different expression pattern (Wilcoxon rank-sum statistic) compared with the old genes (from Figure 10H to Figure 10K), however the gene expression scope becomes border as genes become older. This observation is consistently maintained both for male and female among the two populations. The tissue specific expression illustrates some specialized functions can be performed by young genes, which lead these young genes turn into luxury genes. Usually, the specific expression product in a certain tissue can confer cell-specific structural features and physiological functions, thus promoting the generation of specific phenotypes. Some studies showed that young human genes narrowly and highly express in prefrontal lobes, increasing our human cognition levels compared with other primates, which is an example for the new genes and their specific phenotype (Zhang et al. 2011). Additionally, due to their narrow expression, such gene can be used as feature to evaluate the drug side effects (Duffy et al. 2020). The large-scale census of gene expression scope during gene evolution provides us a new insight into understanding of these phenomena. In our previous section, we modeled the dynamic process of directional gene traffics between the X chromosomes and autosomes based on testis and whole body. The correlations between each pair of biological calculated for the two tissues show high correlations (more than 0.9, Figure 10D), which further demonstrated the high quality of this data and the reliability of our previous result.

### The RNA-based duplication in the *D. melenogaster* subgenus also detected an excess gene traffics from X chromosome to autosomes

Previous analyses revealed a gene traffic out of the X chromosomes in the *Sophophora* subgenus (Betran et al, 2002; Vibranovski et al, 2009). We re-investigated this phenomenon in this subgenus based on our newly dated age. We employed two rounds of BLASTP searches to identify genes origination by RNA-based duplication. In the alignment of young genes (queries) to the old genes (subjects), we selected the best-scoring match, and then it was BLASTed as the query to the young genes (subjects) again. We collected reciprocal best hits (RBH) after the two rounds of BLASTP and annotated young genes were created through RNA-based duplication by their corresponding old genes without introns (Supplementary Figure S7 and Supplementary Table S3_C).

According to previous expectation (Betran et al. 2002), we calculated the expected number of genes moving among chromosomes. In equations (6), (7) and (8),*N_i_* represents the gene number on chromosome *i*, *f*(*N*_*i*_) represents the scaled gene number on chromosome *i*. *L*_*i*_represents the length of chromosome *i*, *f*(*L*_*i*_) represents the scaled length of chromosome *i*. Based on equation 6-8 (Betran et al. 2002), the gene number and chromosome size of a specific chromosome were scaled to [0, 1] according to the total number of protein coding genes and total chromosome size, respectively.

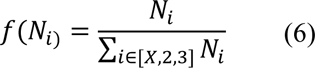

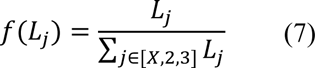

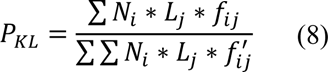

Accordingly, we have the following values: the scaled gene number on chromosome X *N*_*X*_=0.1716, the scaled gene number on chromosome 2 (2L and 2R) *N*_2_=0.3916, the scaled gene number on chromosome 3 (3L and 3R) *N*_3_=0.4368; and also have the scaled chromosome size of chromosome X *L*_*X*_=0.1776, the scaled chromosome size of chromosome 2 *L*_2_=0.3682, the scaled chromosome size of chromosome 3 *L*_3_=0.4541.

According to the pre-computed parameters we detected a significant excess of RNA-based duplicated genes that moved from the X chromosome to an autosomes (X→A) compared with the excepted number (5.9775) and the observed one (13). We also observed a paucity of RNA-based gene movement from autosome to autosome. Both observations are consistent with previous findings in the *Sophophora* subgenus (Betran et al. 2002; Vibranovski et al. 2009), supporting a conclusion that the gene traffic out of the X is a general phenomenon.

## Discussions

### The high-quality genomes provide more precise identification of new genes

We identified a slightly lower percentage of young genes compared with previous databases using earlier genome sequencing techniques. Un-sequenced genomic fragments of lower-quality assemblies can result in absence of alignment between focal and outgroup species, therefore leading to the overestimation of the proportion of young genes (Supplementary Figure S8). In our work we included 9 high-quality PacBio-based assemblies of species in the *Sophophora* group in which these newly included genomes have more than 100.0x coverage allowing us to capture more genome segments in contrast with previous short-read sequencing data (Figure 2A and Figure 2B). For example, the previously sequenced *S. lebanonensis* has 26.8x genome coverage. The species *D. virilis* assembly used by previous studies has 80x genome coverage, which is a much lower genome coverage than that of the assembly we used. Additionally, our incorporation of the evolutionarily most related outgroup species, *S. lebanonensis,* not only avoided overestimating the number of young genes, but to also probe how many gene births occurred before 65 Myr ago when the common ancestors of *S. lebanonensis* and *Drosophila* diverged.

### New genes originated rapidly in two *Drosophila* subgeneras

Gene birth in the *Drosophila* subgenus had not been previously explored because too few outgroup species in the subgenus had been sequenced which would have resulted in low resolution. By adding more species belonging to the *Drosophila* subgenus and including newly sequenced species of the *Sophophora* subgenus we were able to study with detail the origination of genes in the *Drosophila* subgenus while avoiding overestimating the number of new genes. By comparing the protein-coding gene births of the two subgenera we observed the rate of protein-coding gene birth in the *Drosophila* and *Sophophora* subgenera to be approximately the same. In contrast with the previous works (Zhou et al. 2008b; Zhang et al. 2010; Shao et al. 2019), our work contained more branches and more high-quality genomes. Therefore, our work improves the resolution of gene age distribution in both subgenera while simultaneously avoiding overestimation of the number of new genes in *D. melanogaster* and *D. virilis*.

### The most recent common ancestor of the *Drosophila* genus acquired rapidly a big number of new genes

The most recent common ancestral species (MRCA) of the *Drosophila* genus stayed in evolution for ∼64 Million years ago before it diverged to two subgenera ∼63 million years ago. In this ancestral stage, acquired ∼900 new protein-coding genes and hundreds of noncoding RNAs. We note here that this high number of new protein-coding genes in a short period is a high rate of new gene evolution. This rate is 4-5 times other metazoans which are often in the range of 15-25 new genes per million years (Long et al, 2013) and several times the new gene evolution in other branches (Figure 3). This species had extensive changes at protein coding level and noncoding RNA level in the genome during its speciation or subsequent evolution in a short period.

These changes might lead to extensive divergence of subgenera and rapid radiation to ∼ 2,000 species as known today in the genus (O’Grady and DeSalle 2018).

**Table 1.**
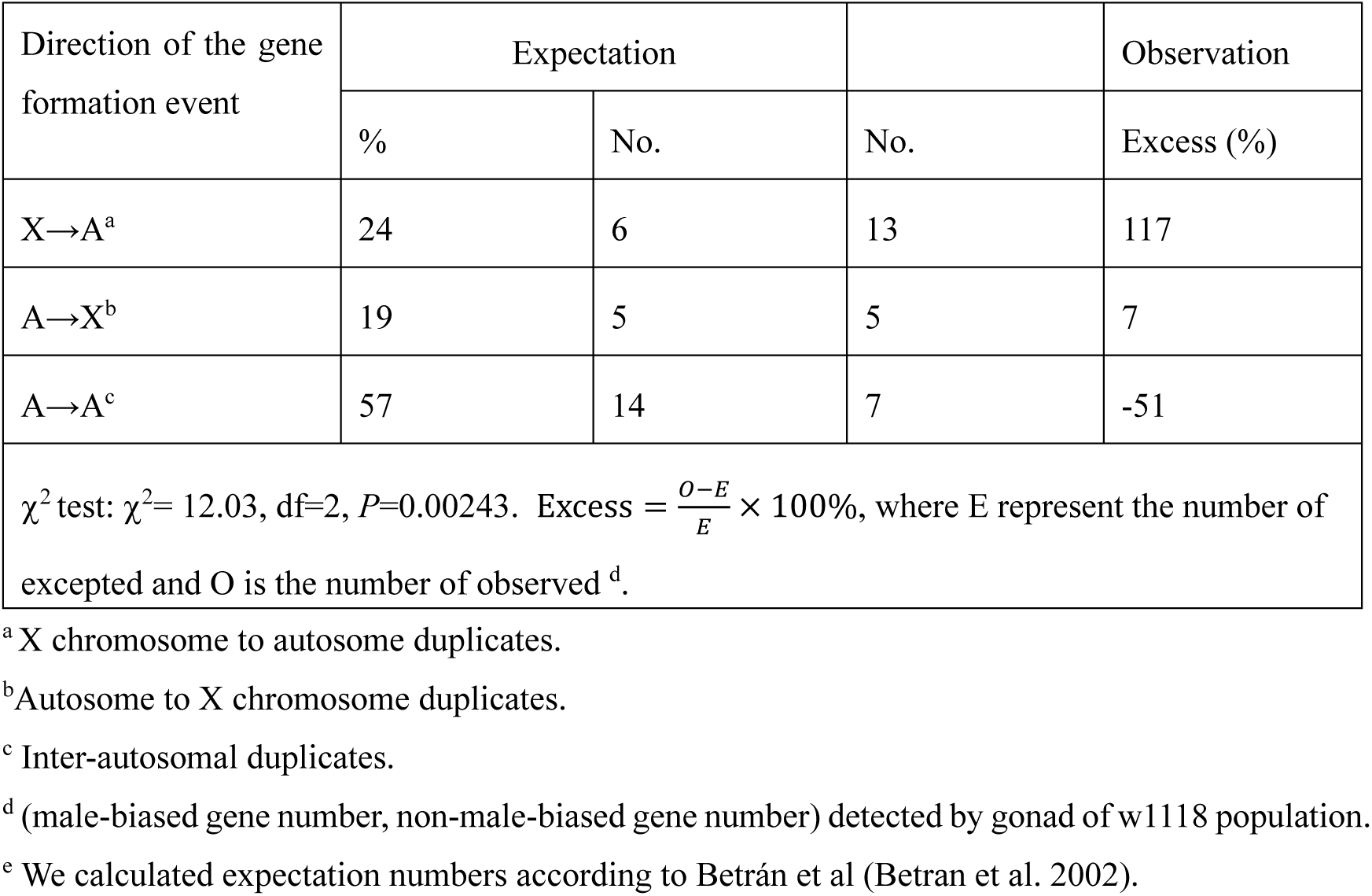
RNA-based movement pattern

## Supplemental Materials

Supplementary File S1: Evolutionary tree in a Newick format for dating gene age of *D. melenogaster*

Supplementary File S2: Evolutionary tree in a Newick format for all avaiable *Drosophila* species both for NCBI and TimeTree

Supplemental File S3: Evolutionary tree in newick format for dating gene age of *D. virilis*

Supplementary Table S1: *Drosophila* species listed in NCBI. Supplementary Table S2: Genome assemblies list in dating gene age

Supplementary Table S3_A: Protein-coding gene age list of *D. melenogaster*

Supplementary Table S3_B: ncRNA gene age list of *D. melenogaster*

Supplementary Table S3_C: Retroposed new genes in *D. melanogaster*

Supplementary Table S4: Gene age list of *D. virilis*

Supplementary Table S5: Gene expression fold change based on gonad tissur of w1118

Supplementary Table S6: Gene expression fold change based on gonad tissur of orgR

Supplementary Table S7: Gene expression fold change based on whole body expresson of w1118

Supplementary Table S8: Gene expression fold change based on whole body of orgR

Supplementary Figure S1. Density estimation for confidence score provided in COMPARTMENTS database

Supplementary Figure S2. Correlation analysis between EI and species divergent time

Supplementary Figure S3. The distribution of sex-biased genes among young and old genes, which is inferred based on expression data in whole body and two *D. melanogaster* populations.

Supplementary Figure S4. The sex-biased gene identification based on gonad and the correlation analysis

Supplementary Figure S5. The sex-biased gene identification based on whole body and the correlation analysis

Supplementary Figure S6. The modelling of male-based genes based on whole body expression from two *D. melanogaster* populations during evolutionary time

Supplementary Figure S7. The retro-genes inferred by RBH-based strategy

Supplementary Figure S8. A schematic to explain why sequencing quality can result in overestimating the number of young genes

## Acknowledgement

This study was supported by the National Science Foundation grant (NSF1026200) to M.L., and the scholarship from China Scholarship Council (CSC) to C.D.. We are thankful for the valuable discussion with Chunyan Chen and the members in Long lab. We are also thankful for Ms Wang help us draw the subcellular compartments. We are thankful for the kind discussion about RNA-Seq data analysis with Chengchi Fang and Jianhai Chen.

